# Unified meta regression models for rare variant association studies

**DOI:** 10.1101/2025.01.23.634522

**Authors:** Larissa Lauer, Manuel A. Rivas

## Abstract

Rare variant association studies (RVAS) of complex traits have emerged as a powerful approach to advance drug discovery and diagnostics. Missense pathogenicity predictions from AlphaMissense based on structural context and protein language models improve the differentiation between benign and deleterious variants. Constraint metrics, on the other hand, allow researchers to pinpoint genomic regions under selective pressure that may not directly impact protein structure, but are more likely to contain functionally important mutations. Loss-of-function (LoF) variants, which result in the complete or partial loss of protein function, are particularly informative, as it is more straightforward to assess their downstream functional consequences. In this study, we present a unified meta regression model approach that incorporates the probability of pathogenicity, probability of constraint, and indicator whether a variant is a predicted loss-of-function or missense variant as features to model the observed effect size and uncertainty of effect size obtained from single-variant genetic analysis. We applied the unified meta regression model to 1,144 continuous phenotypes from UK Biobank using single variant summary statistics obtained from Genebass. We replicated our findings using the AllofUS cohort. For each gene discovery, we make available a characterization of whether constrained sites are associated with the phenotype, whether pathogenic sites determined by structural based predictions are associated with phenotype, and whether broader loss-of-function or missense variant annotation better explains the summary statistics observed. Our results are publicly available at Global Biobank Engine (https://biobankengine.shinyapps.io/phenome-wide-unified-model/).

## I. INTRODUCTION

Genome-wide association studies (GWAS) have successfully mapped over a thousand regions of the genome to more than a hundred human traits^1^. These studies typically test for association between approximately a million genetic variants and a single trait (e.g. disease or biomarker measurement)^2^. For the past decade due to technological constraints the majority of variants tested have been common in the population, i.e. with a frequency of at least 5% in the general population. With the data generation capacity of high-throughput sequencing technologies, it has become possible to assess the relevance of rare variants. Henceforth, we refer to studies of common variants as *Common Variant Association Studies* (CVAS), and studies of rare variants as *Rare Variant Association Studies* (RVAS).

In rare variant association studies, the methodological considerations differ from common variant association studies^3^. First, we typically need to consider how to improve power to detect the association between rare variants and a trait^4^. One strategy is to aggregate the signals of associations across a *testing unit*, like a protein-coding gene^5^. Then, within the testing unit, restrict the association analysis to variants with a particular annotation class like *protein-truncating variants*^*6,7*^, which are variants predicted to introduce a premature stop when the gene is translated to a protein. Finally, after testing for association we obtain a p-value for the unit tested (e.g. gene).

Recent studies have found that there are locations within the genome that are constrained, i.e. mutations are observed in those locations with lower probability compared to the rest of the genome^8,9^. Initial studies have found that constrained genes are more likely to be associated with rare diseases and reduced reproductive success^10,11^. Additionally, researchers have identified enrichment in association between constrained regions of the genome and neuropsychiatric conditions like schizophrenia^12^.

We can view a rare variant association analysis within a testing unit as a special case of a meta regression analysis. In meta regression analysis we fit the observed effect size estimates from univariate regression to the potential *moderators* in the testing unit, e.g. annotation of a variant like protein-truncating variant or missense variant, or other quantitative measurements of pathogenicity of the variant like a log transformation of the probability of constraint^13^, or a log transformation of the probability of pathogenicity obtained from protein-folding prediction algorithms like Google DeepMind’s AlphaMissense scores^14^, or a measure of evolutionary constraint as obtained using the PrimateAI algorithm^15^. Another potential moderator is the readout from massively parallel reporter assays, which can provide a continuous measurement of the effect of a genetic variant on gene expression^16^. Overall, meta regression modeling provides a flexible and efficient framework where data across multiple fronts can be combined to improve **power to detect and explain associations in rare variant association studies**.

Here, we present a unified meta regression model that combines data from four moderators with summary level data from univariate regressions applied to 1,144 phenotypes from 394,841 individuals from the UK Biobank. We present gene-based association statistics and moderator level p-values along with an estimate of the proportion of variance explained by moderators where gene-based association was detected (p *<* 1*×*10^−6^, see **Figure 1**).

**FIG. 1:**
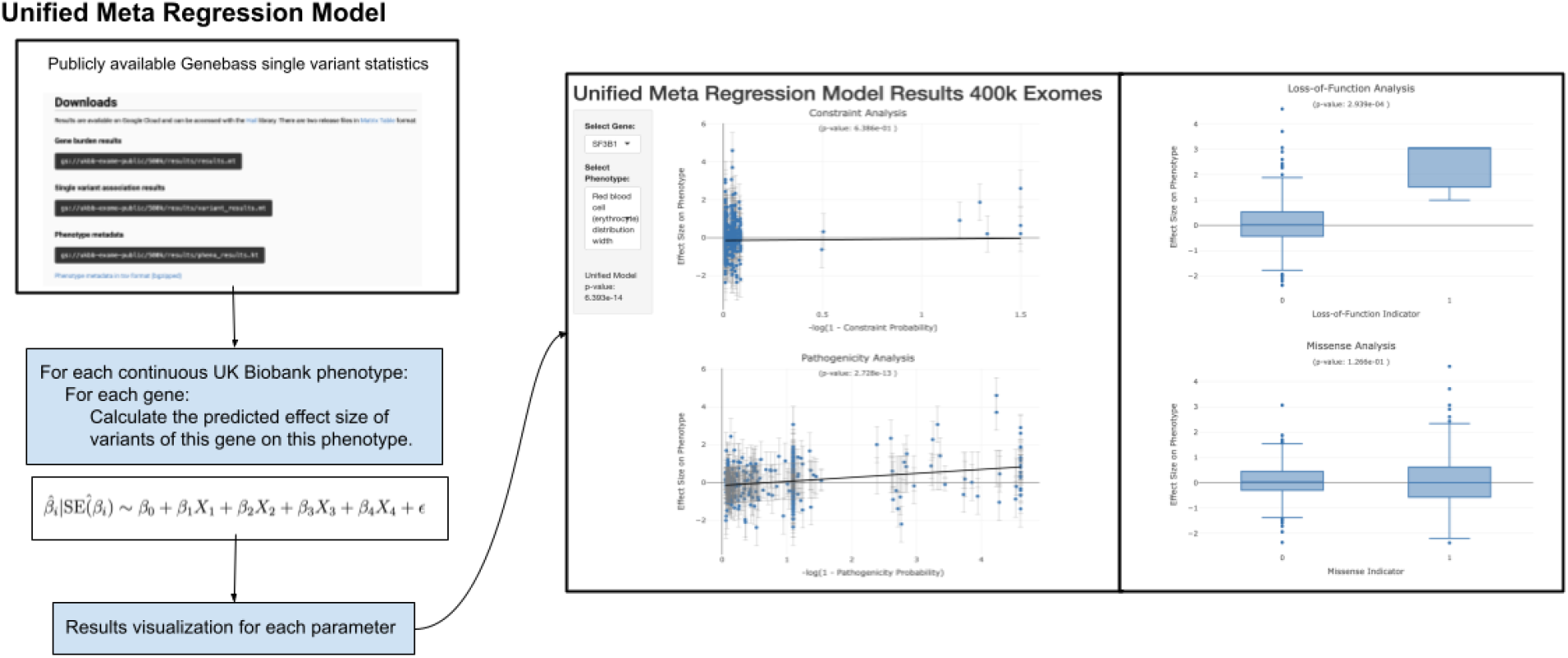
Flowchart for unified meta regression models. Summary statistics are obtained from large-scale exome sequencing studies like those available in Genebass.org. Unified meta regression models are applied to univariate summary statistic data and integrated with moderator data like constraint probabilities and pathogenicity probabilities from AlphaMissense. Visualization of the results from the meta regression models are supported in Global Biobank Engine.

## II. METHODS

Assume we have 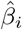 and standard error (SE),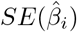, estimates from a univariate regression model applied to a rare variant *i*, for variants *i* = 1, …, *m* in a gene *k*.

### A. Model setup

The meta-regression model is:

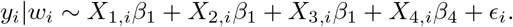

The weights are 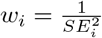. The weighted least squares coefficient estimates are given by:

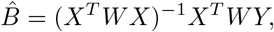

where *X* is the design matrix of size *m × p*, where *m* is the number of variants in the gene and *p* corresponds to the number of predictors/moderators.

Here, we restrict *p* to be indexed by *j* for each variant *i* the following moderators:

- *X*_*i*,1_ = − log(1 − probability of constraint_*i*_), which is obtained from the Hidden Markov Model;
- *X*_*i*,2_ = − log(1 − probability of pathogenicity_*i*_), which is obtained from AlphaMissense;
- *X*_*i*,3_ = **1**_*LoF,i*_, which is an indicator variable whether the variant is a Loss-of-function variant (= 1) or not (= 0); and
- *X*_*i*,4_ = **1**_*Missense,i*_, which is an indicator variable whether the variant is a Missense variant (= 1) or not (= 0).
- *W* is the diagonal weight matrix where *w*_*i*_ is 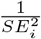.
- *Y* is the outcome vector, which is a vector 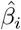, i.e. the regression estimate from the univariate regression applied to the genetic variant data for variant *i* = 1, …, *m*.

### B. Proportion of variance explained by moderators

We can compute the proportion of variance explained by each moderator for the meta-regression model as follows:

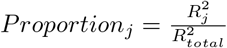for each moderator *j*, where

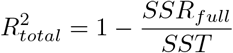

and *SST* is the total sum of squares defined as

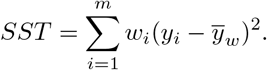

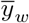 is the weighted mean of *y*:

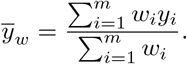

For SSR we can compute

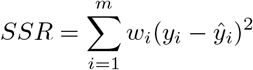

with all moderators (*SSR*_*full*_) and for each moderator by removing the moderator (*SSR*_*reduced,j*_), whereby then we can calculate 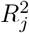,

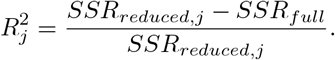

### C. Hypothesis testing for unified model and moderators

The t-statistic for the moderators is given by 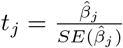, and the moderator p-value is *p*_*j*_ = 2*P* (*t >* |*t*_*j*_|). The F-statistic for the overall model is 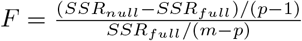, and overall p-value is *p*_*unified*_ = *P* (*F > F*_*observed*_).

## III. RESULTS

We asked how many new gene based association results (p *<* 1 *×* 10^−6^) were obtained using the unified meta regression model. We found 5,134 gene and phenotype combined associations. We identified 867 gene and phenotype combinations with evidence of constraint probability being a significant moderator (p *<* .001), and 2,799 gene and phenotype combinations with evidence of AlphaMissense probability being a significant moderator. 1,045 gene and phenotype combinations have no evidence of the indicator moderator being significant (p*>* .01 for both the indicator variable for missense variant or loss-of-function annotation) for a total of 198 unique protein coding genes where the unified model is exome-wide significant and a combination of the probability moderators are found to be nominally significant. These results highlight the added value of moderators to RVAS of complex traits.

For Lymphocyte count we found 11 genes where constraint or pathogenicity significantly associated with the phenotype (p *<* .001) out of a total of *49* gene-based rare variant associations. We found that constraint probability in *PLEK* is associated with lower lymphocyte counts (see **Figure 2**). *PLEK* encodes Pleckstrin-1, which is a protein that is a member of the Pleckstrins family of proteins found in platelets, which are involved in rearranging the actin cytoskeletion in processes like platelet activation and erythropoeisis.

**FIG. 2:**
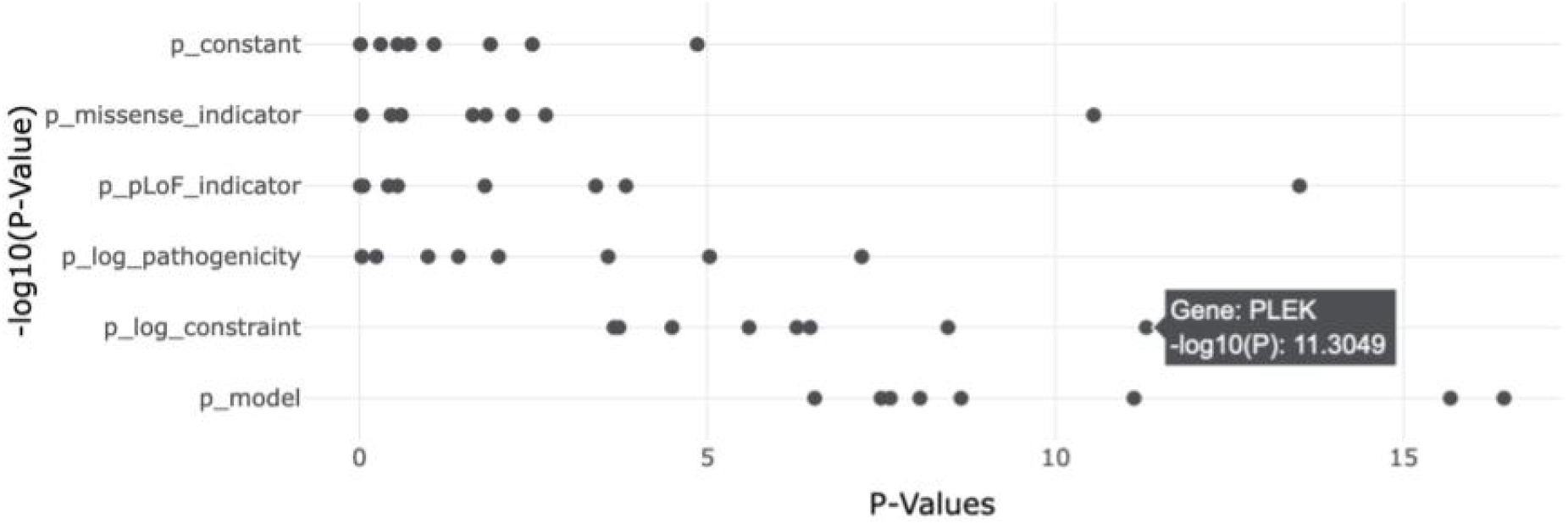
Lymphocyte count meta regression model moderator and unified model p-values. The − log_10_ *P* values (x-axis) for genes associated (unified model [*p*_model_] *p <* 1 *×* 10^6^) with Lymphocyte count and evidence of constraint association [*p*_log constraint_] *p <* .001. *PLEK* is highlighted as having evidence of association between constraint and lymphocyte count phenotype.

For pulse rate and *CASZ1* we found that loss-of-function (*p* = 0.0014), missense (*p* = 4.6 *×* 10^−4^), and constraint probability (*p* = 2.78 *×* 10^−5^) are significantly associated moderators with a unified *p* = 5.28 *×* 10^−13^. *CASZ1* encodes a zinc finger transcription factor that plays a critical role in mammalian cardiac morphogenesis and development. Here, we find that genetic variants that are more likely to be constrained are associated with increased pulse rate (see **Figure 3**).

**FIG. 3:**
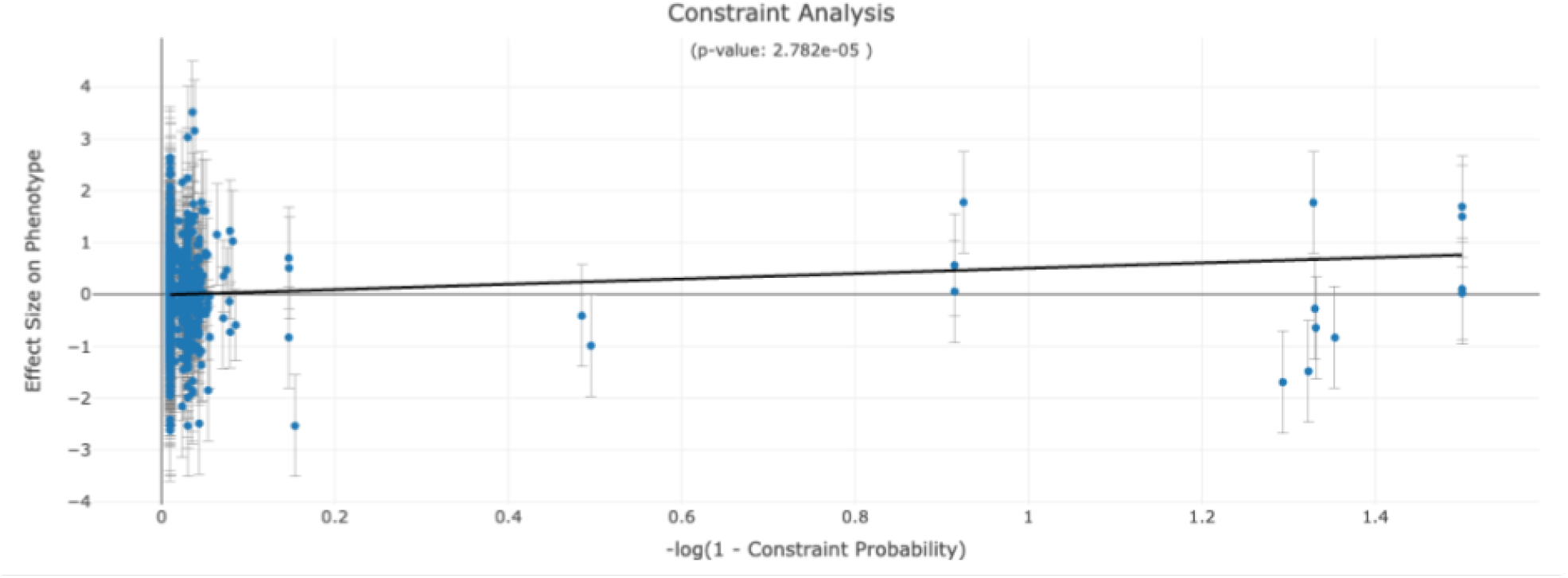
Association between constraint probability and pulse rate. The relationship between the log constraint probability (x-axis) and pulse rate is shown. The estimated effect size for the variant (y-axis) and the standard error estimate (whiskers) are represented. Best fit line is shown. *p* value for the moderator is also shown in the title.

Structural based predictions of pathogenicity aid in assessing whether structure is associated to a phenotype. Here, we found that *SF3B1* is associated to red blood cell (erythrocyte) count (*p* = 2.068 *×* 10^−13^). Although nominal evidence of association (*p <* 0.05) is found for the three moderators: loss-of-function, missense, and constraint probability, strong evidence of association (*p* = 1.60*×*10^−12^) is found for the moderator pathogenicity probability (see **Figure 4**). This indicates that structural based predictions are informative for establishing a relationship between *SF3B1* and red blood cell count. *SF3B1* encodes subunit 1 of the splicing factor 3b protein complex. Mutations in this gene have been recurrently seen in cases of advanced chronic lymphocytic lekeumia, myelodyplastic syndromes and breast cancer.

**FIG. 4:**
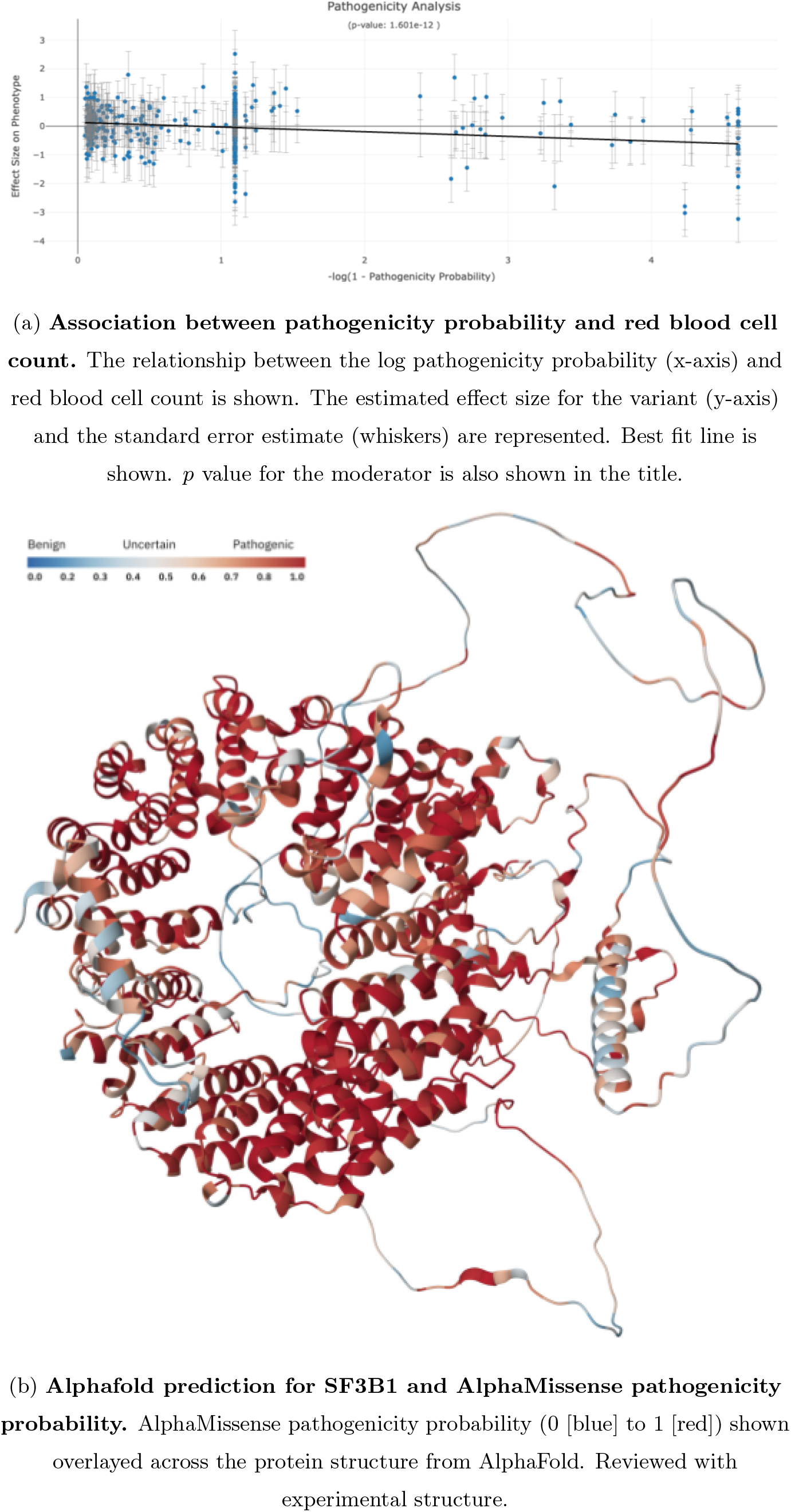
Pathogenicity probability as a moderator in the unified meta regression model.

## IV. DISCUSSION

Our framework for *Rare Variant Association Studies* (RVAS) exploits concepts in meta regression analysis. Here, we propose that modeling summary statistics obtained from univariate regression analysis that is typical of common variant association studies where age, sex, and principal components are included as additional covariates provides a flexible and powerful framework for combining information across multiple variants and multiple moderators.

Moderators, or potentially powerful predictors, of the observed effects between genetic variants and phenotypes could begin to explain why we are seeing signals in rare variant association studies.

Here, we propose four simple moderators: i) whether a variant is a loss-of-function or not, ii) whether a variant is a missense variant or not, iii) the predicted probability of constraint from a simple Hidden-Markov-Model (HMM) learned from millions of exomes^13^, and iv) the predicted probability of pathogenicity obtained from an artificial intelligence model using predicted protein structures.

Analogous to common variant association studies where very powerful studies were achieved by combining summary statistics. Here, for RVAS, we envision that very powerful studies can be conducted by not only sharing summary statistics from univariate regressions, but also sharing of data from potential moderators across the close to a billion variants that will be identified as we go from exome sequencing studies to whole genome sequencing studies.

Previously, we presented efficient regression computation models for rare variant analysis that reduced the time from conducting an exome sequencing study from 695 minutes to 1.6 minutes on a single machine^17^. Using the summary statistics from those studies we can improve inference of rare variants across multiple studies, variants, and moderators. We anticipate that the unified meta regression models framework presented will dramatically improve our ability to conduct and interpret rare variant association studies.

## V. ACKNOWLEDGEMENTS

M.A.R. is in part supported by National Human Genome Research Institute (NHGRI) under award R01HG010140, and by the National Institutes of Mental Health (NIMH) under award R01MH124244 both of the National Institutes of Health (NIH).

The content is solely the responsibility of the authors and does not necessarily represent the official views of the funding agencies; funders had no role in study design, data collection and analysis, decision to publish, or preparation of the manuscript.

